## Supplemental Text for "Algorithms for the selection of fluorescent reporters"

### Supplementary materials for:
### Algorithms for the selection of fluorescent reporters

Prashant Vaidyanathan, Evan Appleton, David Tran, Alexander Vahid, George Church, and Douglas Densmore

### S1 Algorithm Overview

#### S1.1 Prediction of fluorescent signals in a detector

Emission and excitation spectra along with brightness of fluorescent proteins and dyes were obtained from various online databases and repositories [1, 2, 3, 4]. The spectral data for fluorophores was organized into CSV (comma-separated values) file (additional details can be found in sections S2.1.2 and S2.1.3). Each fluorophore has an excitation spectrum and emission spectrum. The exitation spectrum of a fluorophore $f$ can be denoted as: $EX_f : \lambda \rightarrow x$ and the emission spectrum of a fluorophore $f$ can be denoted as: $EM_f : \lambda \rightarrow m$. where $m$ and $x$ are the relative spectral intensities (normalized to 1) and $\lambda$ is the wavelength with values typically ranging from 250nm to 800nm. To compute the amount of fluorophore signal measured by a detector, we consider the wavelength of the laser ($\lambda_l$) associated with the detector ($d$). We then look-up the excitation value ($\chi = EX_f(\lambda_l)$) associated with the wavelength in the excitation spectrum of the fluorophore. This excitation value determines the intensity of fluorescence that can be observed in the detector. We then scale the Emission Spectrum of the fluorophore by $\chi$:

$$EM_f^{\lambda_l}(\lambda) = (\chi) * EM_f(\lambda) \tag{S1}$$

If we also consider the brightness of the fluorophore ($\beta_f$), this brightness value is normalized to 1 based on the brightness of the brightest fluorophore in our library (Section S2.1.3). We then scale the Emission Spectrum of the fluorophore with the product of $\beta$ and $\chi$ as follows:

$$EM_f^{\lambda_l}(\lambda) = (\chi * \beta) * EM_f(\lambda) \tag{S2}$$

We compute the area under the curve within the length of the detector and divide it by the width of the detector. We approximate the area under the curve using Left Riemann Sum (Section S1.2).

#### S1.2 Left Riemann Sum

Reimann sum is a way of approximating the area under a curve. It is an approximation of an integral by a finite sum. We have chosen the left Reimann sum method to compute the amount of signal that is captured by a detector. Let's assume that we wish to find the signal captured by a detector $d$ for a fluorophore $f$ which is excited by a laser $l$ of wavelength $\lambda_l$. Let $[\lambda_{start}, \lambda_{end}]$ denote the range of wavelengths covered by $d$. We first create a partition $P$ of $[\lambda_{start}, \lambda_{end}]$:

$$P = [x_0, x_1], [x_1, x_2], ..., [x_{n-1}, x_n] \tag{S3}$$

where $\lambda_{start} = x_0 < x_1 < x_2 < ... < x_{n-1} < x_n = \lambda_{end}$ and all subintervals are of equal length. The left Reimann sum is given by:

$$S = \sum_{i=1}^{n} EM_f^{\lambda_l}(x_{i-1}) * (x_i - x_{i-1}) \tag{S4}$$

where $EM_f^{\lambda_l}$ is obtained from Eq.S1 (or Eq.S2 if we consider brightness as well). We then normalize the value to 1 by dividing $S$ by $(\lambda_{end} - \lambda_{start})$ (i.e. the width of the detector).

#### S1.3 Computational prediction of signal and bleed-through values in a panel

We use the left reimann sum to calculate the signal and bleed-through values in each detector of the panel. This is done by using Eq. S4 for the wavelength of the laser associated with the detector. Table S1 shows an example of the signal and bleed-through values in each detector of a 5-color fluorophore panel designed for the CytoFlex. The diagonal contains the signal values (marked in bold). These values are displayed by the fpSelection tool.

While comparing the computational predictions with experimental measurements, the values in each detector are normalized to the highest value in that detector. In an ideal panel, the diagonal (signal) values will be 1, and the bleed-through values will be 0 (or close to 0). In an invalid panel, one or more signal values (diagonal values) will be zero. Table S2 shows the normalized values from Table S1.

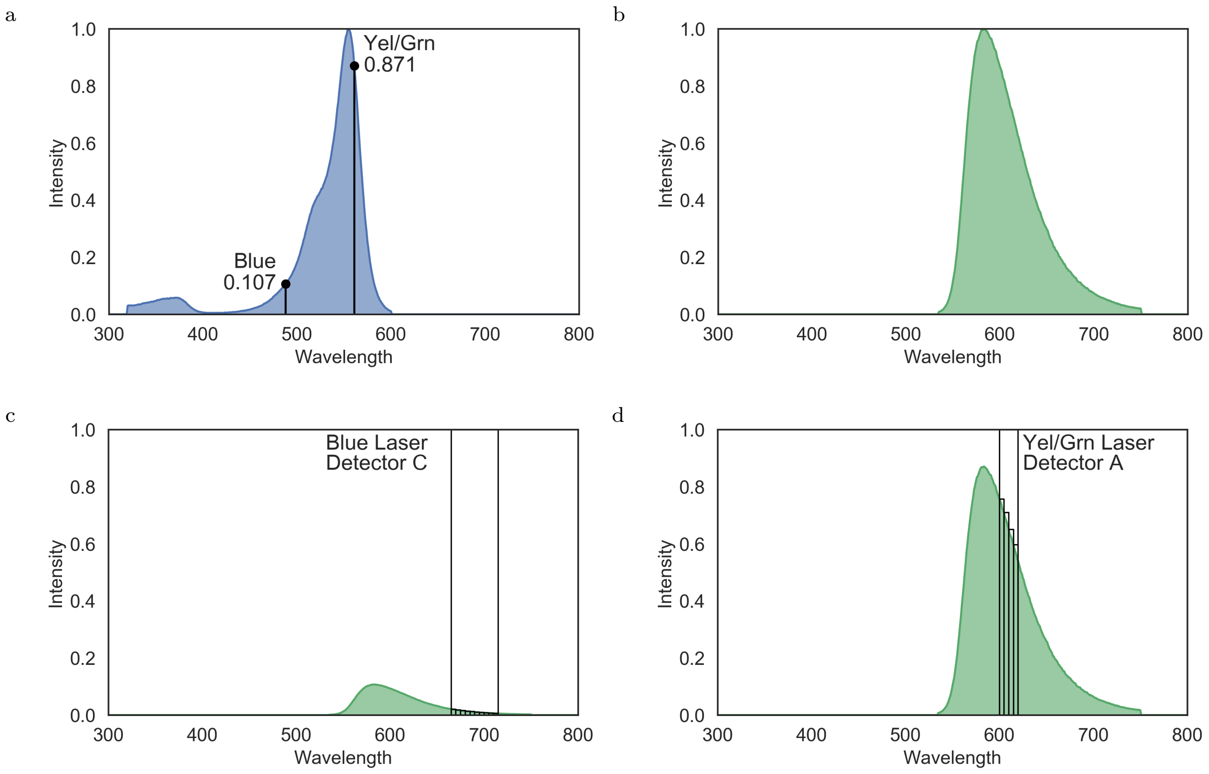

Figure S1: Predicting TagRFP signal in detectors of a CytoFlex LX cytometer. S1a shows the excitation spectrum of TagRFP while S1b shows the peak emission spectrum of TagRFP. S1c shows the emission of TagRFP when excited by a Blue laser at 488nm. To calculate signal detected by Detector C (690/50 BP), the reimman left sum is computed by creating a partition from 665 to 715 (with subinterval length 5) and using eq.S4 which yields 0.60281, which when normalized to 1 (by dividing the sum by the length of the detector = 50) gives 0.01205 or 1.205%. Similarly, S1d shows the emission of TagRFP when excited by a Yel/Grn laser at 561nm. The reimman left sum under Detector A (610/20 BP) is 13.5721 which when normalized to 1 gives 0.67860 or 67.860%.

| | **Violet-B** | **UV-A** | **Red-C** | **Yel/Grn-C** | **Yel/Grn-D** |
|---|---|---|---|---|---|
| **Cerulean** | **0.29402** | 0 | 1.26E-06 | 0.000332 | 4.48E-05 |
| **Sirius** | 0.00423 | **0.63** | 0 | 0 | 0 |
| **iRFP713** | 0 | 0 | **0.50014** | 0 | 0.009438 |
| **tdTomato** | 0.00012 | 0 | 0 | **0.798275** | 0.138683 |
| **mPlum** | 0 | 0 | 0 | 0.063856 | **0.523409** |

Table S1: Computational prediction of Signal and Bleed-through values in a 5-color fluorophore panel.

| | **Violet-B** | **UV-A** | **Red-C** | **Yel/Grn-C** | **Yel/Grn-D** |
|---|---|---|---|---|---|
| **Cerulean** | **1** | 0 | 4.29E-06 | 0.001129 | 0.000152 |
| **Sirius** | 0.006681 | **1** | 0 | 0 | 0 |
| **iRFP713** | 0 | 0 | **1** | 0 | 0.01887 |
| **tdTomato** | 0.000154 | 0 | 0 | **1** | 0.173728 |
| **mPlum** | 0 | 0 | 0 | 0.121999 | **1** |

Table S2: Signal and Bleed-through values normalized to the brightest value in each detector.

#### S1.4 Search Algorithms

We developed three algorithms to explore the search space of n-color cytometry panels for a given set of fluorophores and instrument configuration - Exhaustive Search, Hill Climbing, and Simulated Annealing. The algorithms use the method described in the Methods section to compare two n-color panels to determine which configuration is more optimal or desirable. Exhaustive search uses a brute force technique by exploring the entire search space to find the optimal solution among valid panel designs. Hill climbing and simulated annealing start with a random n-color panel and randomly swap either a detector or a fluorophore to incrementally find better solutions. These two algorithms are heuristics that don't always guarantee optimal results (especially for larger values of $n$) but have negligible run times (compared to the exhaustive algorithm). Hence these algorithms can be run multiple times to obtain better results.

As described in the main text, the optimality of a design is determined by two factors:

- Validity of the panel. A valid n-color panel is one where all the detectors in the panel have a non-zero value of signal of the fluorophore it has been assigned to detect (irrespective of the amount of bleed-though from other fluorophores in the panel).
- If both panels are valid, then we consider the following factors in this order:
  - The number of detectors where bleed-through is within some specified threshold ($\eta$).
  - Geometric mean of the predicted signal values in all the detectors.
  - Arithmetic mean of the predicted overall bleed-through values in all the detectors.

The search algorithms first simulate the amount of signal and bleed-through expected in each detector of the panel and use the factors listed above to compare the optimality of the panel with another panel.

Since simulated annealing consistently finds good solutions and has a fast run-time, it is the default algorithm used in the web application. However, users can use the other algorithms via the command line application.

##### S1.4.1 Exhaustive Search

The exhaustive search algorithm is a brute-force search that looks for all possible combinations fluorophore panels of size $n$. Algorithm 1 shows the pseudo-code for exhaustive search. For a library of fluorophores $F$ and set of detectors $D$ of a measurement instrument, the total number of n-color fluorophore panels are:

$$\frac{|F|!}{(|F|-n)!} * \frac{|D|!}{(|D|-n)!} * \frac{1}{n!} \tag{S5}$$

The solution space is the Cartesian product of all permutations of $n$ fluorophores and combinations of $n$ detectors (or vice-versa). However, not all solutions in this space are valid. The exhaustive search algorithm can be optimized with a few preprocessing steps to avoid in-valid solutions. For instance, certain lasers may not excite a specific fluorophore (which can be determined by observing the normalized excitation spectrum values corresponding to the wavelength of the laser). Hence, all panels containing a detectors of lasers that do not excite the fluorophore can be disregarded. Similarly, if a fluorophore's normalized emission spectrum value is 0 for all wavelengths covered by a detector, then all panels where that detector has been assigned to the fluorophore can be disregarded.

Additional computational optimizations such as multi-threading can also be used to speed up the exhaustive search. However, as shown in Fig. S2, the exhaustive search algorithm does not scale for larger values of $n$ or for larger fluorophores libraries and larger number of detectors. For instance, exhaustively searching for all 5-color fluorophore panels from a library of 12 fluorophores for the CytoFlex which has 19 detectors, can take more than 2 hours to run, while searching for an optimal 6-color fluorophore panel for the same instrument is estimated to take more than 48 hours to run.

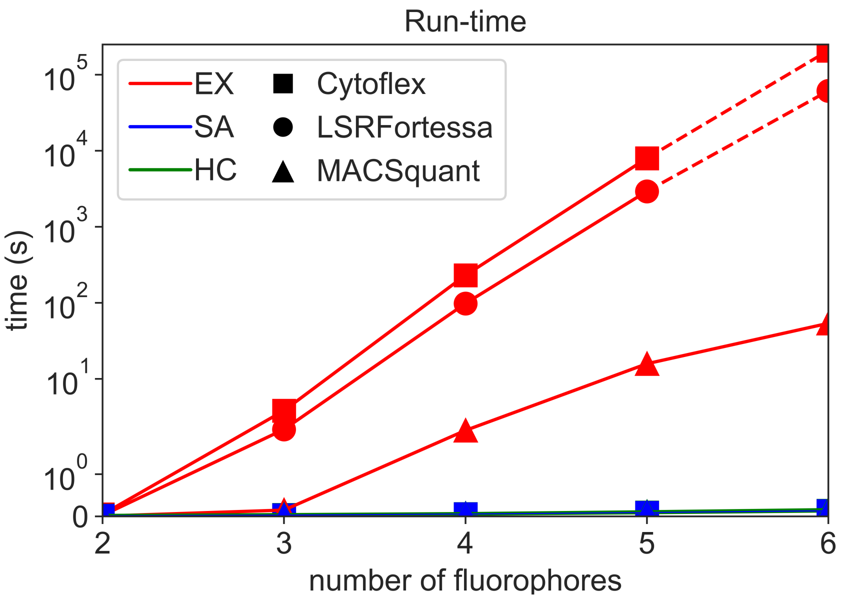

Figure S2: Run-time for Exhaustive (EX), Hill Climbing (HC), and Simulated Annealing (SA), find n-color panels from a library of 12 fluorophores. The CytoFlex instrument had 19 detectors to choose from, where as the LSRFortessa had 16, and the MACSquant had 7 detectors. The x-axis shows the size of the panel while the y-axis shows time in seconds. It is quite clear that the run-time for Exhaustive search has exponential growth and becomes intractable for higher values of n. The dotted line shows the predicted time required to run exhaustive search for n=6 for the LSRFortessa and CytoFlex instruments based on the average amount of time required to evaluate a single 6-color panel and compare it to another 6-color panel.

##### S1.4.2 Hill Climbing

The hill climbing algorithm is a heuristic which incrementally tries to find a better solution. The algorithm starts with an arbitrary n-color panel by randomly choosing $n$ distinct fluorophores and detectors. The algorithm then compares neighboring panels by randomly swapping either a fluorophore or detector to the current solution. If a neighboring panel is better than the current solution, it swaps to the neighbor as the current best solution. This is done for 10000 iterations and the final best panel is returned as the solution. If the final panel is not valid, the algorithm starts over from a new arbitrary starting point, until the algorithm times out. Algorithm 2 shows the pseudocode for the Hill Climbing algorithm.

The challenge of choosing a valid, optimal n-color panel is not a convex optimization problem. Hence, the hill climbing approach often returns a local optima as the result as shown in Fig.S3. While hill climbing is marginally faster than simulated annealing, this speed gain in run-time is negligible since both algorithms take less than 1 second to complete a run. Since the results of simulated annealing are consistently better, we recommend simulated annealing over hill climbing.

##### S1.4.3 Simulated Annealing

The simulated annealing algorithm uses a probabilistic technique to find a global optimum. The algorithm starts with an "annealing temperature" which gradually reduces via a cooling rate. Like hill climbing, simulated annealing starts with an arbitrary n-color panel by randomly choosing $n$ unique fluorophores and detectors. At each iteration, of the run, the algorithm computes an acceptance probability switching to a neighboring panel (which like hill climbing is obtained by randomly swapping either a fluorophore or a detector). The acceptance probability is determined based on the properties of the current as well as the neighboring panel: the number of detectors with non-zero signal (x), the number of detectors where the overall bleed-through is within the specified threshold $\eta$ (y), the geometric of the predicted signal values in the detectors (z), and the arithmetic mean of the predicted overall bleed-through values in the detectors (w).

**Algorithm 1** Exhaustive Search

**Input:** List of fluorophores $F$, List of detectors $D$, Size of panel $n$
**Output:** A Panel $(F_n, D_n)$ of size $n$.

fluorophorePermutations $\leftarrow$ Generate all permutations of $n$ fluorophores out of $F$
fluorophorePermutations $\leftarrow$ Generate all combinations of $n$ detectors out of $D$
$F_{best} \leftarrow$ fluorophorePermutations.get(0)
$D_{best} \leftarrow$ detectorCombinations.get(0)
**for each** f **in** fluorophorePermutations **do**
    **for each** d **in** detectorCombinations **do**
        **if** $(f, d)$ is not valid **then**
            **continue**
        **end if**
        **if** $(f, d)$ is better than $(F_{best}, D_{best})$ **then**
            $(F_{best}, D_{best}) \leftarrow (f, d)$
        **end if**
    **end for**
**end for**
**return** $(F_{best}, D_{best})$

**Algorithm 2** Hill Climbing

**Input:** List of fluorophores $F$, List of detectors $D$, Size of panel $n$
**Output:** A Panel $(F_n, D_n)$ of size $n$.

iterations $\leftarrow$ 10000
$F_{best} \leftarrow$ Randomly choose $n$ unique fluorophores from $F$
$D_{best} \leftarrow$ Randomly choose $n$ unique detectors from $D$
**for** $i \leftarrow 0$; $i <$iterations; $i \leftarrow i + 1$ **do**
    $(F_{new}, D_{new}) \leftarrow (F_{best}, D_{best})$
    Randomly pick either a fluorophore or detector in $(F_{new}, D_{new})$ and replace it with another fluorophore or detector (respectively). If the new chosen object already exists in $F_{new}$ or $D_{new}$ in another slot, swap the slots of the new chosen object and the object to be swapped.
    **if** $(F_{new}, D_{new})$ is better than $(F_{best}, D_{best})$ **then**
        $(F_{best}, D_{best}) \leftarrow (F_{new}, D_{new})$
    **end if**
**end for**
**return** $(F_{best}, D_{best})$

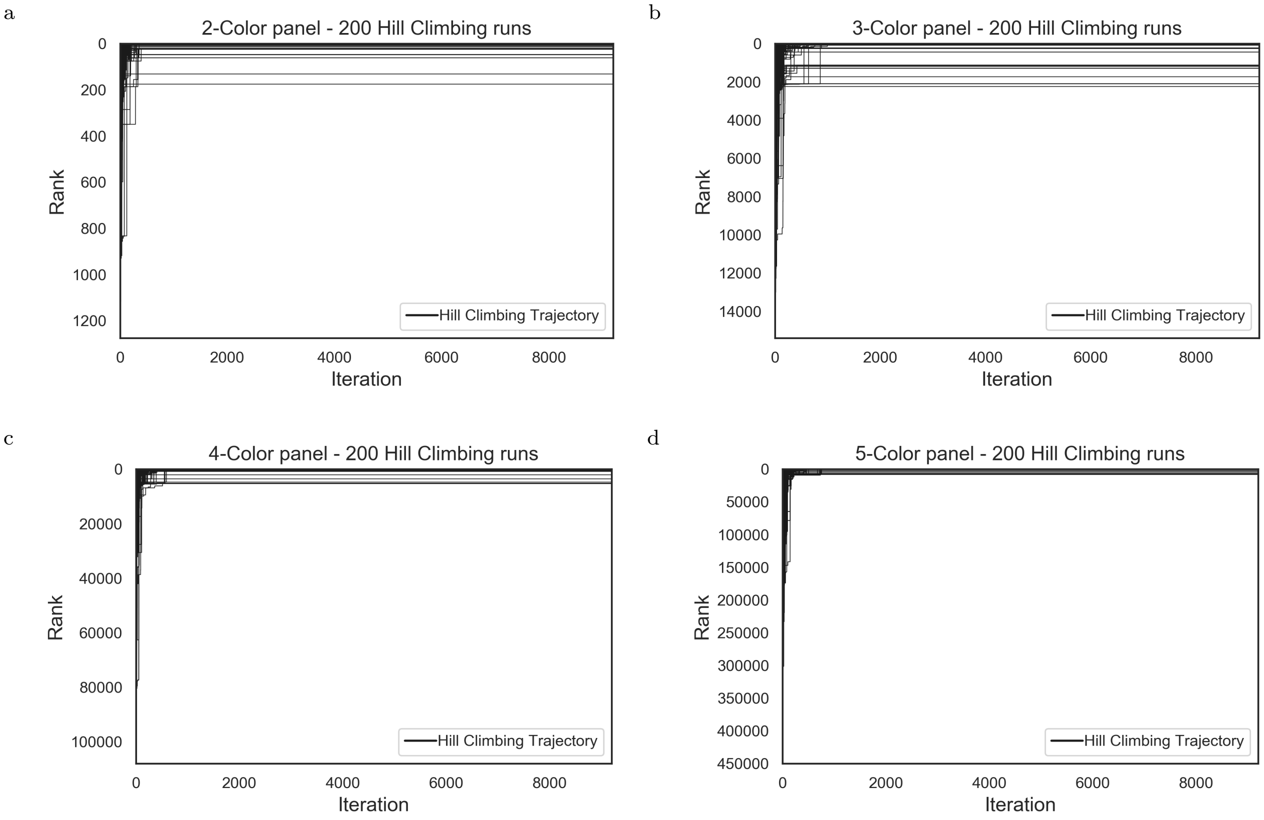

Figure S3: Trajectories of various runs of Hill Climbing. We used exhaustive search to rank all valid solutions for n-color panels for the CytoFlex measurement instrument which has 19 detectors (where rank 0 indicates the best solution). We used a library of 8 fluorophores instead of 12 due to computational memory limitations. We then ran simulated annealing 200 times for values of n ranging from 2 to 5. Each trajectory shows the rank of the best solution at any given iteration of the hill climbing run.

If the neighboring panel has a greater value of x (number of non-zero signals), it is immediately accepted. If both panels have the same value of x, then y values are compared. If the neighboring panel has a greater value of y, then it is immediately accepted. However, if both x and y values are same for the current and the neighboring panel, the acceptance probability is computed based on the z values (or w values if the z values are equal). The probability of accepting a worse solution reduces as the annealing temperature decreases. Algorithm 3 shows the pseudocode for the simulated annealing approach.

In our case studies, we have set the starting temperature as 1000 with a cooling rate of 0.003. Hence, each simulated annealing run has 9206 iterations. Fig.S4 shows the trajectories of 200 runs of simulated annealing for 2,3,4, and 5 color panels for the LSR CytoFlex machine. The web-application runs 50 concurrent threads of simulated annealing runs and returns the best solution among the 50 threads as the result. This is particularly useful for higher values of $n$. Alternatively, users can also modify the cooling rate or initial annealing temperature in the source code to achieve better results, especially if the solution space is too large.

Fig.S4 shows the trajectories of 200 runs of simulated annealing for panel sizes ranging from 2 to 5. We can compare the efficiency of Simulated Annealing against Hill Climbing by comparing the results of these runs. For instance, consider the results of simulated annealing and hill climbing for 5-color fluorophore panels (Fig.S4d and Fig.S3d respectively). For a library of 8 fluorophores, and a measurement instrument with 19 detectors, there are 78,140,160 5-color panels of which 449,762 are valid. Simulated annealing was able to find a valid solution in all 200 runs and was able to find the best solution 191 times, whereas hill climbing found a valid solution only 163 times and found the best solution only 75 times. The average rank of the

**Algorithm 3** Simulated Annealing

**Input:** List of fluorophores $F$, List of detectors $D$, Size of panel $n$
**Output:** A Panel $(F_n, D_n)$ of size $n$.

t $\leftarrow$ 1000
coolingRate $\leftarrow$ 0.003
**function** GETACCEPTANCEPROBABILITY($(F_c, D_c)$, $(F_n, D_n)$, temp)
    $(x_c, y_c, z_c, w_c) \leftarrow$ GETPANELPROPERTIES($(F_c, D_c)$)
    $(x_n, y_n, z_n, w_n) \leftarrow$ GETPANELPROPERTIES($(F_n, D_n)$)
    **if** $x_n < x_c$ **then return** 1.0
    **else if** $x_n > x_c$ **then return** 0.0
    **else**
        **if** $y_n > y_c$ **then return** 1.0
        **else if** $y_n < y_c$ **then return** 0.0
        **else**
            **if** $z_c$ **not equals** $z_n$ **then**
                **return** $1.0/(1.0 + e^{-(z_n - z_c)/temp})$
            **else**
                **return** $1.0/(1.0 + e^{-(w_c - w_n)/temp})$
            **end if**
        **end if**
    **end if**
**end function**
$F_{best} \leftarrow$ Randomly choose $n$ unique fluorophores from $F$
$D_{best} \leftarrow$ Randomly choose $n$ unique detectors from $D$
**while** t $> 1$ **do**
    $r \leftarrow$ random number between 0 and 1 (both included).
    $(F_{new}, D_{new}) \leftarrow (F_{best}, D_{best})$
    Randomly pick either a fluorophore or detector in $(F_{new}, D_{new})$ and replace it with another fluorophore or detector (respectively). If the new chosen object already exists in $F_{new}$ or $D_{new}$ in another slot, swap the slots of the new chosen object and the object to be swapped.
    **if** GETACCEPTANCEPROBABILITY($(F_{best}, D_{best})$,$(F_{new}, D_{new})$, t) $> r$ **then**
        $(F_{best}, D_{best}) \leftarrow (F_{new}, D_{new})$
    **end if**
    t $\leftarrow$ t * (1 - coolingRate)
**end while**
**return** $(F_{best}, D_{best})$

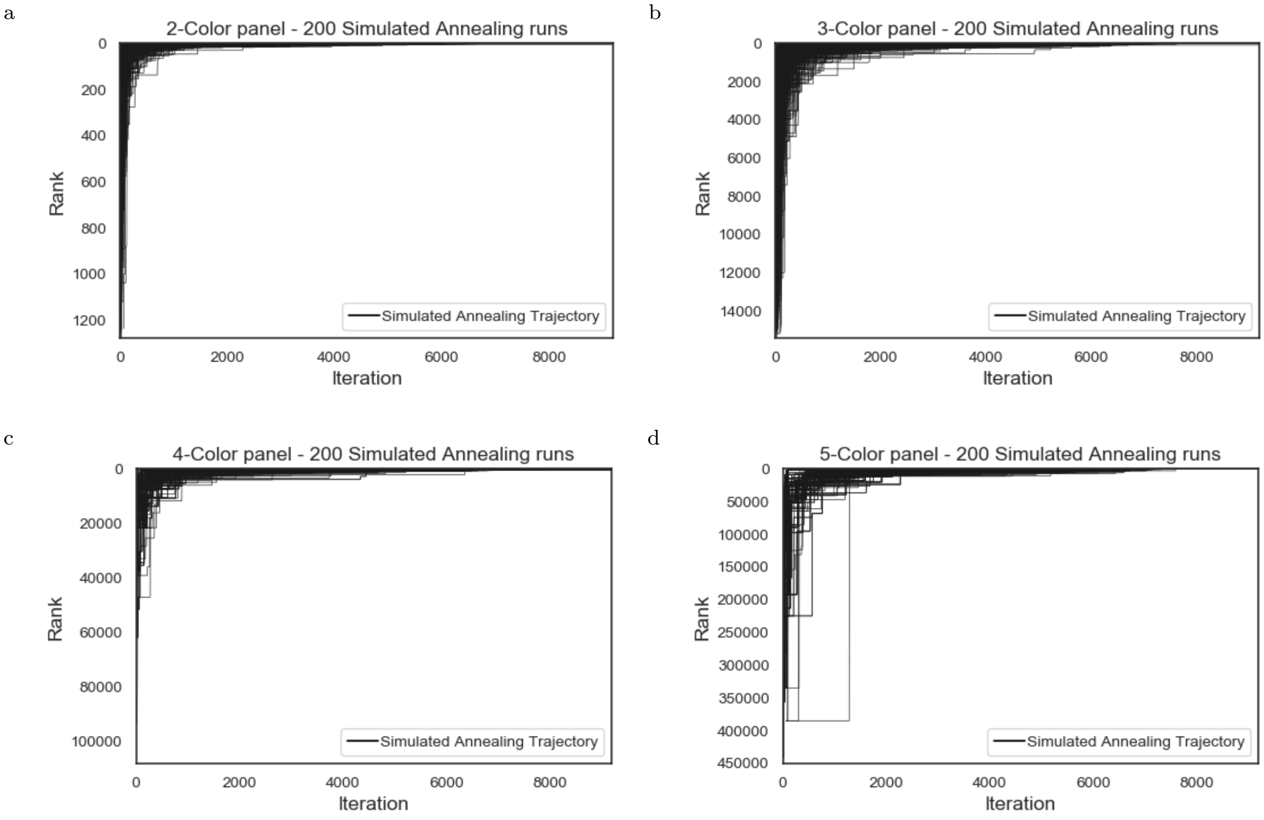

Figure S4: Trajectories of various runs of Simulated Annealing. We used exhaustive search to rank all valid solutions for n-color panels for the CytoFlex measurement instrument which has 19 detectors (where rank 0 indicates the best solution). We used a library of 8 fluorophores instead of 12 due to computational memory limitations. We then ran simulated annealing 200 times for values of n ranging from 2 to 5. Each trajectory shows the rank of the best solution at any given iteration of the simulated annealing run.

solutions found by simulated annealing was 2.005 with a standard deviation of 15.718. Among all the valid solutions found by hill climbing, the average rank of the solutions was 23556.93 with a standard deviation of 57964.5.

### S2 Software Overview

We created a web-application FP Selection (hosted at http://fpselection.org/) which allows users to choose $n$ fluorophores from a given list of fluorophores for a specific cytometer configuration. The code for this software tool is open-source and is hosted at https://github.com/CIDARLAB/fpSelection.

#### S2.1 Inputs

FP Selection, accepts CSV (Comma-Separated Values) files as inputs. Sample files can be found in the "FPSelection.zip" file. FP Selection also has additional tools so that users can generate input files using manual data entry (http://fpselection.org/tools).

##### S2.1.1 Instrument configuration

We adapted the format of the configuration file generated by the BD LSRFortessa cytometer. Table S3 shows an example of an instrument configuration CSV file. (The configuration files of the instruments used

| Laser Name | Type | Wavelength | Power | Detector Array | Detector | Channel | Mirror | Filter | Parameter | Fsc Channel | Position |
|---|---|---|---|---|---|---|---|---|---|---|---|
| Blue | Custom | 488 | | Custom | A | 1 | | 525/40 BP | B525 FitC | | 1 |
| | | | | | B | 2 | | 610/20 BP | B610 ECD | | |
| | | | | | C | 3 | | 690/50 BP | B690 PC5 | | |
| Red | Custom | 640 | | Custom | A | 4 | | 763/43 BP | R763 APC750 | | 4 |
| | | | | | B | 5 | | 660/10 BP | R660 APC | | |

Table S3: Instrument configuration file

in the case study can be found in "FPSelection.zip").

The parser extracts the name of the laser, the laser wavelength, list of detectors, detector names, and filter mid-point and width from the configuration file. The table shown below has additional meta-data (such as laser power, laser position, etc) which is stored in the Laser and Detector data-structures, however, these properties are not currently used to design fluorophores.

##### S2.1.2 Fluorophore spectra

The emission and excitation spectra is specified via a CSV file (the file used in the case study can be found in "FPSelection.zip"). Table S4 shows the format for the spectra file. The CSV must contain a column with a header: **Wavelength -** which contains wavelengths specified as a floating point values. Each fluorophore must have two columns, one for excitation where the column header is specified as **FluorophoreName (EX)** and one for emission with column header formatted as **FluorophoreName (EM)**. The spectra file can also be used to specify the emission and excitation spectra of the autofluorescence of the cell-line. The algorithm treats this like any other fluorophore and assigns a detector to detect autofluorescence and factors the fluorescence emission of the cell as bleed-through in other detectors of the panel.

| **Wavelength** | **Cerulean (EX)** | **Cerulean (EM)** | **KO (EX)** | **KO (EM)** |
|---|---|---|---|---|
| 540 | 0.0036 | 0.3783 | 0.7962 | 0.1269 |
| 541 | 0.0036 | 0.3723 | 0.8275 | 0.1493 |
| 542 | 0.0047 | 0.3622 | 0.8637 | 0.1768 |
| 543 | 0.0042 | 0.354 | 0.8972 | 0.2088 |
| 544 | 0.0046 | 0.346 | 0.9286 | 0.2424 |
| 545 | 0.0046 | 0.339 | 0.9585 | 0.2846 |
| 546 | 0.0047 | 0.3308 | 0.9756 | 0.3317 |
| 547 | 0.0053 | 0.3161 | 0.9909 | 0.3825 |
| 548 | 0.0048 | 0.3106 | 0.9998 | 0.4386 |
| 549 | 0.0055 | 0.3004 | 1.0000 | 0.5012 |
| 550 | 0.0046 | 0.2942 | 0.9913 | 0.5613 |

Table S4: Fluorophore Spectra format

##### S2.1.3 Brightness

Users have the option to upload the brightness of the fluorophores via a CSV file (the file used in the case study can be found in "FPSelection.zip"). The brightness file is not required and if it is not specified, the brightness of all fluorophores is considered equal. The CSV contains just two columns, the first is the name of the fluorophore, while the second is the brightness value (an example of this is shown in table S5). The brightness values are normalized to 1 based on the brightness fluorophore in the library. The normalized brightness value ($\beta$) is used in Eq.S2. The fluorophore names in the first column must match the names in the spectra CSV file.

For the fluorophores in table S5, the brightness is normalized to the brightness of tdTomato (the brightest fluorophore in the list). Hence the normalized brightness values are 1.0 for tdTomato, 0.060491 for iRFP720, and 0.73514 for mScarlet.

| Fluorophore | Brightness |
|---|---|
| tdTomato | 95.22 |
| iRFP720 | 5.76 |
| mScarlet | 70 |

Table S5: Brightness file

It is important to note that the tool currently considers brightness of fluorophores as a relative value. Hence, if the brightness considered is the molecular brightness (or theoretical brightness computed by multiplying the extinction coefficient and quantum yield), then this brightness must be used for all fluorophores in the library to get consistent results. It is also important to remember that various sources of fluorophores scale and report brightness based on distinct scales or parameters. For instance, the brightness of dyes is often reported on a scale of 1 to 5 where 1 is the dimmest and 5 is the brightest, which is different compared to the molecular brightness specified in FPbase (like the values shown in Table S5).

#### S2.2 Graphical User Interface

The web-application for fpSelection is hosted at http://fpselection.org/. Fig.S5 shows the main page of the tool. Users can upload instrument settings, fluorophore spectra, fluorophore brightness and select the size of the fluorophore panel they wish to design. By clicking the **GO!** button, fpselection runs 50 concurrent threads of simulated annealing to find the optimal solution.

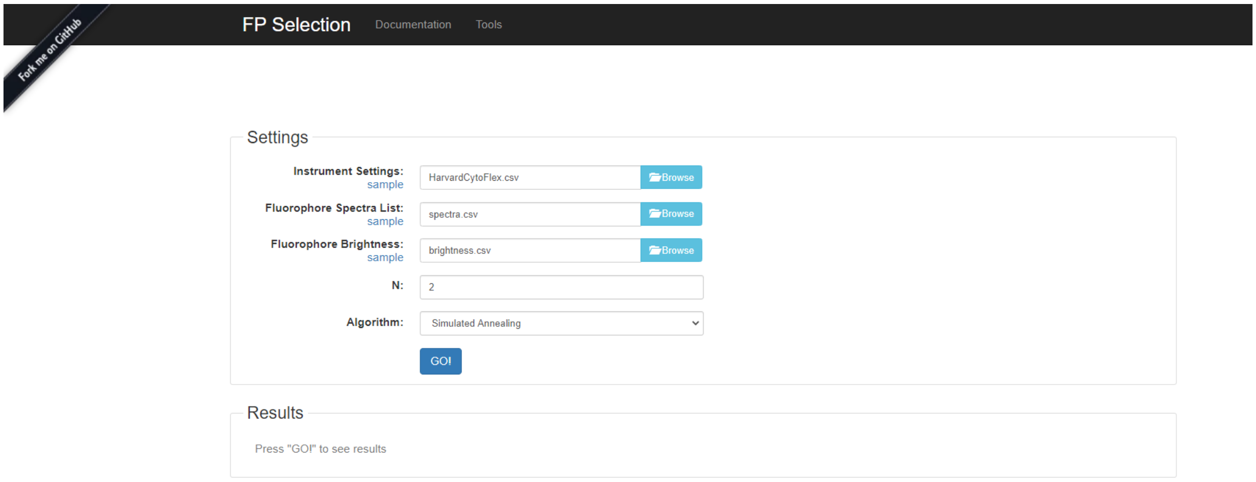

Figure S5: Main Page of fpselection.org

##### S2.2.1 Understanding the Results

The results are displayed as plots to visualize the predicted signal and bleed-through in each detector of the panel. In each subplot, the x-axis shows the wavelength in nm, the y-axis shows the relative intensity of emission normalized to 1. The rectangular plot shows the detector and the range of wavelengths it covers. The signal is shown in green, while the sum of all bleed-through from all other fluorophores in the panel is shown in red. The web-app also displays the panel information in text format by listing the laser, detector, and the fluorophore assigned to the detector. Fig.S6 shows a screenshot of the results produced by the webapp for a 2-color panel.

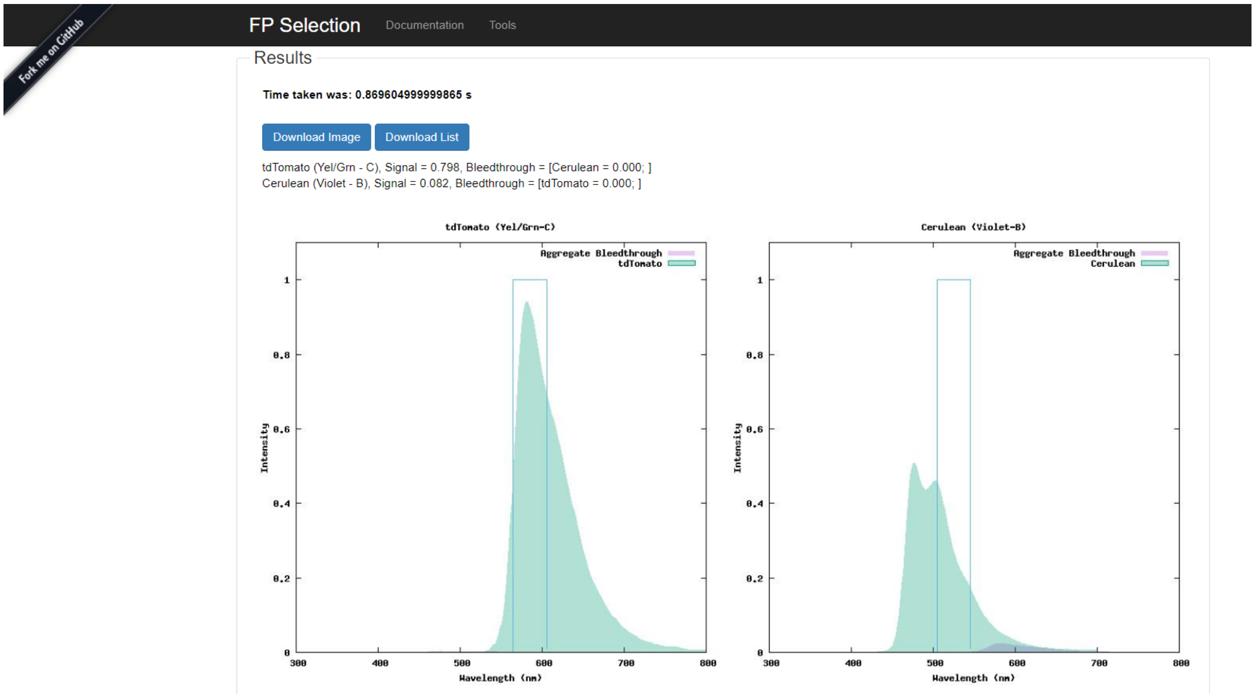

Figure S6: Result shown by fpselection.org for 2-color fluorophore panel for the CytoFlex cytometer where fluorophores were chosen from a library of 12 fluorophores.

##### S2.3 Technology

The code for fpSelection was written in Java using the Apache Maven framework. The web-app is built using Jetty framework. The tool is capable of plotting using both matplotlib (which requires Python) and JavaPlot (which requires gnuplot). The source code is hosted on GitHub: https://github.com/CIDARLAB/fpSelection, where users can find instructions to build and test the command-line tool and can also deploy a local version of the web-app. The tool has been tested in both Windows 10 as well as Ubuntu 18.04 LTS operating systems.

#### S3 Fluorophore library

Table S6 lists the 12 fluorophores that were used in the case study. For computational runs where 8 fluorophores were used, we used all fluorophores listed except tdTomato, mScarlet, mCherry, and KO.

The emission and excitation spectra were obtained primarily from FPbase[1]. We used the molecular brightness listed in FPbase. However, as specified in previous sections, this brightness value does not always reflect what is observed experimentally. The full emission and excitation spectra can be found in "FPSelection.zip".

| Fluorophore | Theoretical Brightness | Peak Emission (nm) | Peak Excitation (nm) |
|---|---|---|---|
| mCherry | 15.84 | 587 | 610 |
| mOrange | 48.99 | 548 | 562 |
| DsRed2 | 28.6 | 578.5 | 593.5 |
| iRFP713 | 6.3 | 689 | 713 |
| Sirius | 3.6 | 354 | 425 |
| Cerulean | 26.66 | 435 | 477 |
| KO | 49.39 | 549 | 561 |
| mPlum | 4.1 | 587 | 650 |
| TagRFP | 48 | 554 | 584 |
| tdTomato | 95.22 | 556 | 581.5 |
| iRFP720 | 5.76 | 701 | 720 |
| mScarlet | 70 | 570 | 594 |

Table S6: Fluorophore Library

### S4 Measurement Instruments

Table S7 lists the lasers and detectors of the CytoFlex LX cytometer, while table S8 lists the configurations of the BD LSRFortessa cytometer and table S9 lists the configurations of the Miltenyi MACSquant VYB cytometer. In all these cytometers, the detectors were photomultiplier tubes (PMTs). The tables list the excitation wavelengths of the lasers (in nanometers) and the mid point and filter length of the bandpass filters associated with the PMTs (in nanometers).

| Laser | Wavelength | Detector | Filter (Midpoint/width) |
|---|---|---|---|
| Blue | 488 | A | 525/40 BP |
| | | B | 610/20 BP |
| | | C | 690/50 BP |
| Red | 640 | A | 763/43 BP |
| | | B | 660/10 BP |
| | | C | 712/25 BP |
| Violet | 405 | A | 450/50 BP |
| | | B | 525/40 BP |
| | | C | 610/20 BP |
| | | D | 660/10 BP |
| | | E | 763/43 BP |
| UV | 355 | A | 405/30 BP |
| | | B | 525/40 BP |
| | | C | 675/30 BP |
| Yel/Grn | 561 | A | 610/20 BP |
| | | B | 763/43 BP |
| | | C | 585/42 BP |
| | | D | 675/30 BP |
| | | E | 710/50 BP |

Table S7: CytoFlex LX Lasers and Detectors

| **Laser** | **Wavelength** | **Detector** | **Filter (Midpoint/width)** |
|---|---|---|---|
| Blue | 488 | A | 685/35 BP |
| | | B | 540/25 BP |
| | | C | 515/20 BP |
| Red | 640 | A | 780/60 BP |
| | | B | 670/30 BP |
| Violet | 405 | B | 582/15 BP |
| | | C | 525/50 BP |
| | | D | 515/20 BP |
| | | E | 450/50 BP |
| UV | 355 | A | 530/30 BP |
| | | B | 450/50 BP |
| Yel/Grn | 561 | A | 780/60 BP |
| | | B | 710/50 BP |
| | | C | 660/20 BP |
| | | D | 610/20 BP |
| | | E | 582/15 BP |

Table S8: BD LSRFortessa Lasers and Detectors

| **Laser** | **Wavelength (nm)** | **Detector** | **Filter (Midpoint/width)** |
|---|---|---|---|
| Violet | 405 | A | 450/50 BP |
| | | B | 525/50 BP |
| Blue | 488 | A | 525/50 BP |
| | | B | 614/50 BP |
| Yel/Grn | 561 | A | 586/15 BP |
| | | B | 615/20 BP |
| | | C | 661/20 BP |

Table S9: Miltenyi MACSquant VYB Lasers and Detectors
